# Identification of novel H2A histone variants across diverse clades of algae

**DOI:** 10.1101/2024.06.14.598977

**Authors:** Ellyn Rousselot, Zofia Nehr, Jean-Marc Aury, France Denoeud, J Mark Cock, Leïla Tirichine, Céline Duc

## Abstract

Histones are recognized as one of the most conserved proteins across eukaryotes. They not only ensure DNA compaction in the nucleus but also participate in epigenetic regulation of genome expression. These key epigenetic players are divided into canonical histones, expressed during the S-phase, and variants, expressed throughout the cell cycle. Compared with other core histones, H2A proteins exhibit a high level of variability but the characterization of algal H2A variants remains very limited. In this study, we exploited genome data from 21 species to perform a phylogenetic analysis of brown seaweeds. We identified new H2A variants that are specific to the Phaeophyceae and closely-related sister group species. For each identified H2A protein, we characterized motifs present in the histone fold domain as well as in the N- and C-terminal tails. This approach enabled us to classify the different isoforms and to identify three new variants (H2A.N, H2A.O and H2A.E). Based on mass spectrometry and RNA-Seq data, we identified distinct epigenetics marks and specific expression patterns of the H2A variants in both vegetative and reproductive tissues as well as in response to various environmental conditions. These analyses identified sperm-specific variants and inferred putative roles for the newly discovered variants.

## INTRODUCTION

In eukaryotes, DNA is compacted into chromatin within the nucleus. The basic subunit of chromatin is the nucleosome, which is composed of ∼147bp of DNA wrapped around a histone octamer (Luger et al. 1997). The H3-H4 tetramer is associated with two H2A-H2B dimers to compose the histone core. Core histones have amino- and carboxy-terminal tails harbouring various post-translational modifications that contribute to epigenetic information and a characteristic domain known as the histone-fold domain. This histone fold is made of three α-helices (α1, α2, and α3) connected by short loops L1 and L2 (Luger et al. 1997). Histone proteins isoforms can differ in the number of variable amino acids to different extents. Histones are classified into two main categories: (i) canonical histones synthetized during the S-phase and (ii) histone variants or replacement histones expressed throughout the cell cycle. Like the other core histones, H2A participates to nucleosome stability and DNA compaction and has specialized variants. Indeed, most eukaryotes have at least three histone isoforms, canonical H2A plus H2A.X and H2A.Z variants. While the biological functions of canonical H2A remain elusive, H2A variants are actively studied. The H2A.X has a predominant role not only in genome stability as a marker of DNA damage events but also in a wide variety of cellular processes such as chromosome segregation or development (Turinetto and Giachino 2015). The H2A.Z variant is involved in various mechanisms such as transcription regulation, DNA repair, differentiation and is linked to several cancers associated with epigenetic disorders (reviewed in Giaimo et al. 2019; Tsukii et al. 2022).

All H2A isoforms have an acidic patch composed of acidic residues in their C-terminal tails. These H2A acidic residues, together with those of H2B, interact with the H4 N-terminal tail (Luger et al. 1997). In addition, the docking domain, which includes the α3 and αC helices, enables interaction with the H4 C-terminal tail (Luger et al. 1997). The H2A.X variant is characterized by a SQ[D/E] [F/Y] motif at the very end of its C-terminal tail with a serine that is phosphorylated in response to double-strand DNA breaks in animals and yeast (Rogakou et al. 1999) as well as in plants (Lorković et al. 2024). The H2A.Z variant harbours a LEYLTAEVLELAGNA signature in the α2 helix (Jiang et al. 1998) and an L1 loop with a single amino acid insertion, an acidic patch with an extra acidic residue and a docking domain that is one amino acid shorter. Additional H2A variants have been identified in some eukaryotes. For example, vertebrates have a macroH2A protein harbouring a macro-domain in its C-terminal tail and mammals have several additional H2A variants, namely H2A.B, H2A.P and H2A.L. Seed-bearing plants have a characteristic H2A variant, H2A.W, which possesses a stretch of lysine and glycine residues in its N-terminal tail, a RY[A/S][Q/K] sequence in its L1 loop (Kawashima et al. 2015) and a KSPKK C-terminal motif: this variant has a critical role in heterochromatin formation (Yelagandula et al. 2014; Lorković et al. 2017). In addition, land plants other than Angiosperms have another specific variant, H2A.M. This variant, which is present in the liverwort *Marchantia polymorpha* and the moss *P. patens*, is characterized by a RYA[Q/K] sequence in its L1 loop (Kawashima et al. 2015).

Few studies have identified histones in algae (Veluchamy et al. 2015; Bourdareau et al. 2021; Rommelfanger et al. 2021; Khan et al. 2018) consequently leaving an important knowledge gap regarding algal histones. The investigation of histone variants in algae is relatively limited so far due to the considerable diversity within this group, which includes both unicellular and multicellular organisms spanning independent diverse phylogenetic clades, and due to the lack of available genomic and transcriptomic data. There are three main clades of algae (Rhodophyta, Chlorophyta and Chromalveolata). Rhodophyta and Chlorophyta belong to the kingdom Archaeplastida, a major lineage of photosynthetic organisms that also includes land plants. The brown algae belong to the Chromalveolata lineage. Only limited number of genomes have been available for algal species but the recent release of the Phaeoexplorer dataset has provided high quality reference genomes of brown seaweeds (Denoeud et al. 2024). Thanks to this significant advancement, our phylogenetic approach led to an in-depth analysis of H2A evolution in various algal clades and the identification of three novel H2A variants that occur either in several algal lineages or are specific to brown seaweeds. Our results demonstrate that (i) PTM (Post-Translational Modifications) are deposited on a restricted set of H2A variants and (i) there is differential expression of H2A variants during the life cycle of brown seaweeds and in various aquatic habitats. Therefore, this study significantly advances our understanding of histones in eukaryotes.

## RESULTS

### Identification of putative canonical H2A isoforms and the novel H2A.N variant

In our previous study, we identified various histone proteins in brown algae and described H3 isoforms based on sequence similarity with mammalian ones (Denoeud et al. 2024). In order to identify and characterise H2A isoforms in brown algae, we took advantage of a set of 21 reference genomes from the Phaeoexplorer database to identify all the predicted H2A proteins at both the transcript and the gene level (Supplemental_Table_S1). These genomes belong to 17 species from seven different orders of the Phaeophyceae class (thereafter referred as brown seaweeds) and 4 species (*Heterosigma akashiwo, Chrysoparadoxa australica, Schizocladia ischiensis* and *Tribonema minus*) are included as closely-related outgroup species. The chosen species represent different ecological niches located at various levels of the intertidal and subtidal zones, a complex life cycle and various genome sizes. We also added data from *C. tenellus* to have a more comprehensive view of the evolutionary relationship between the brown algal orders Discosporangiales and Dictyotales. We used several well-defined signatures of H2A variants to classify these proteins from brown seaweeds.

Only *H. akashiwo, C. australica* and *T. minus*, which are Phaeophyceae-closely-related sister group species have canonical H2A (Fig. 1A) *i.e.* proteins that do not have the SQ[D/E]Y motif or H2A.Z features (Supplemental_Fig_S1A) and are of similar length and sequence to canonical H2A from animals or plants (Supplemental_Fig_S1B). Phaeophyceae (with the exception of *D. mesarthrocarpum* and *C. tenellus*) possess proteins with a longer N-terminal tail extended by approximately 30 amino-acids (Fig. 1B, Supplemental_Fig_S1B-C) with a PLRP motif (Supplemental_Fig_S1D). Since these H2A variants have a long N-terminal tail, we named them H2A.N, following the histone nomenclature established by (Talbert et al. 2012), and performed a closer inspection of their features compared to canonical H2A. The unusual long N-terminal tail shared no sequence homology with other proteins and included a disordered region according to Interproscan (Paysan-Lafosse et al. 2023). No trustworthy structure was detected for this tail region using AlphaFold (Jumper et al. 2021; Varadi et al. 2022). The αN helix of H2A.N has a QSLRA motif (Fig. 1C) that is strongly divergent from motifs of other H2A types such as canonical H2A ([K/R]SSKA for brown algae in Fig. 1D or RSS[K/R]A in animals and plants in Supplemental_Fig_S1E) but with a conserved helix structure. Moreover, the H2A.N variant has a poorly conserved motif in the L1 loop (Kx[A/T]KLxx where x denotes any residue, Fig. 1E) while canonical H2A harbours a more conserved motif (KxGRY[S/A]x, Fig. 1F). Other features, namely the signature in the α2 helix (Fig. 1G-H), the acidic patch signature (Fig. 1I-J) and the docking domains ending with a VLLPKK sequence and a 40-amino-acid region (Supplemental_Fig_S1A,C), are similar between H2A.N and canonical histones. The C-terminal tails of H2A.N proteins are enriched in lysines and acidic residues (Fig. 1B and Supplemental_Fig_S1C) as in plant and animal canonical H2A (Supplemental_Fig_S1F), while canonical H2A proteins of *H. akashiwo, C. australica* and *T. minus* only show lysine enrichment (Fig. 1A and Supplemental_Fig_S1A).

**Figure 1:**
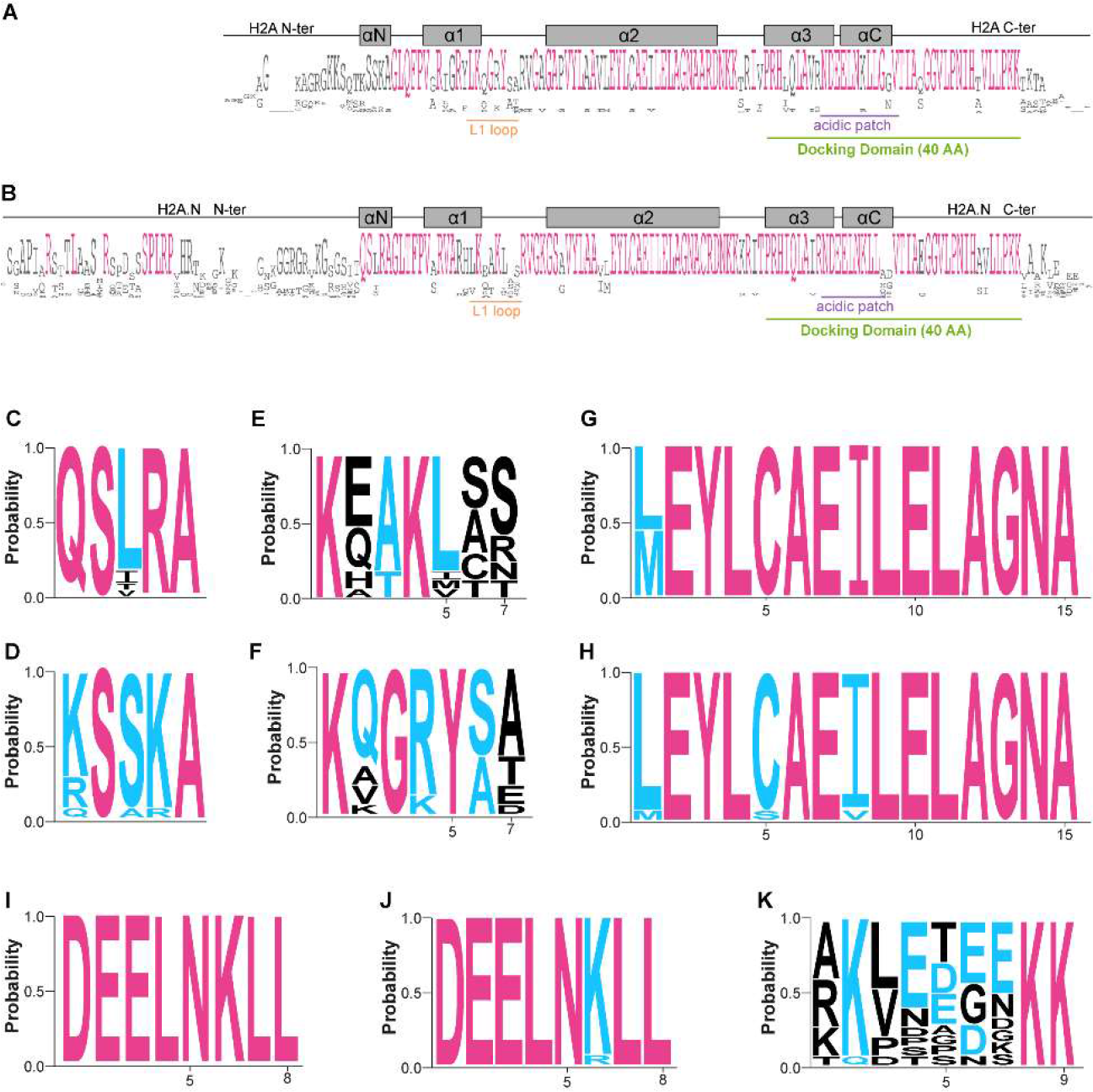
Identification and analysis of canonical H2A and the H2A.N variant in brown algae. *(A-B)* Consensus sequences for canonical H2A (A) and the H2A.N variant sequences (B) generated using Jalview (Waterhouse et al. 2009). The mature protein without the initial methionine is displayed. The L1 loop, acidic patch and docking domain are underlined in orange, purple and green, respectively. The length of the docking domain is indicated (AA, amino acids). The helices are indicated by grey rectangles. N-ter, N-terminal tail; C-ter, C-terminal tail. *(C-D)* Logos of amino acid bias for the αN helix of the H2A.N variant (C) and canonical H2A (D). *(E-F)* Logos of amino acid bias for the L1 loop of the H2A.N variant (E) and canonical H2A (F). *(G-H)* Logos of amino acid bias for the α2 helix of the H2A.N variant (G) and canonical H2A (H). *(I-J)* Logos of amino acid bias for the acidic patch of the H2A.N variant (I) and canonical H2A (J). *(K)* Logo of amino acid bias for C-terminal tail of the H2A.N variant.

### Identification of H2A.X variants in brown seaweeds

We identified H2A.X variants only in the Discosporangiales and the sister outgroup clades of the Phaeophyceae (Raphidophyceae, Chrysoparadoxophyceae, Xanthophyceae, Schizocladiophyceae). Identification was based on the presence of a SQ[D/E]Y motif at the end of the C-terminal tail and the absence of a H2A.Z signature (Fig. 2A, Supplemental_Fig_S2). These proteins have a KQG[R/Q]Y[S/A] [T/A/K] motif in their L1 loop (Fig. 2B), a [L/M]EYLTAEVLELAGNA degenerate signature in their α2 helix (Fig. 2C) and a DEELNKLL motif in the acidic patch (Fig. 2D). The H2A.X docking domain ends with a VLLPKK sequence and is 40 amino acids long (Fig. 2A, Supplemental_Fig_S2).

**Figure 2:**
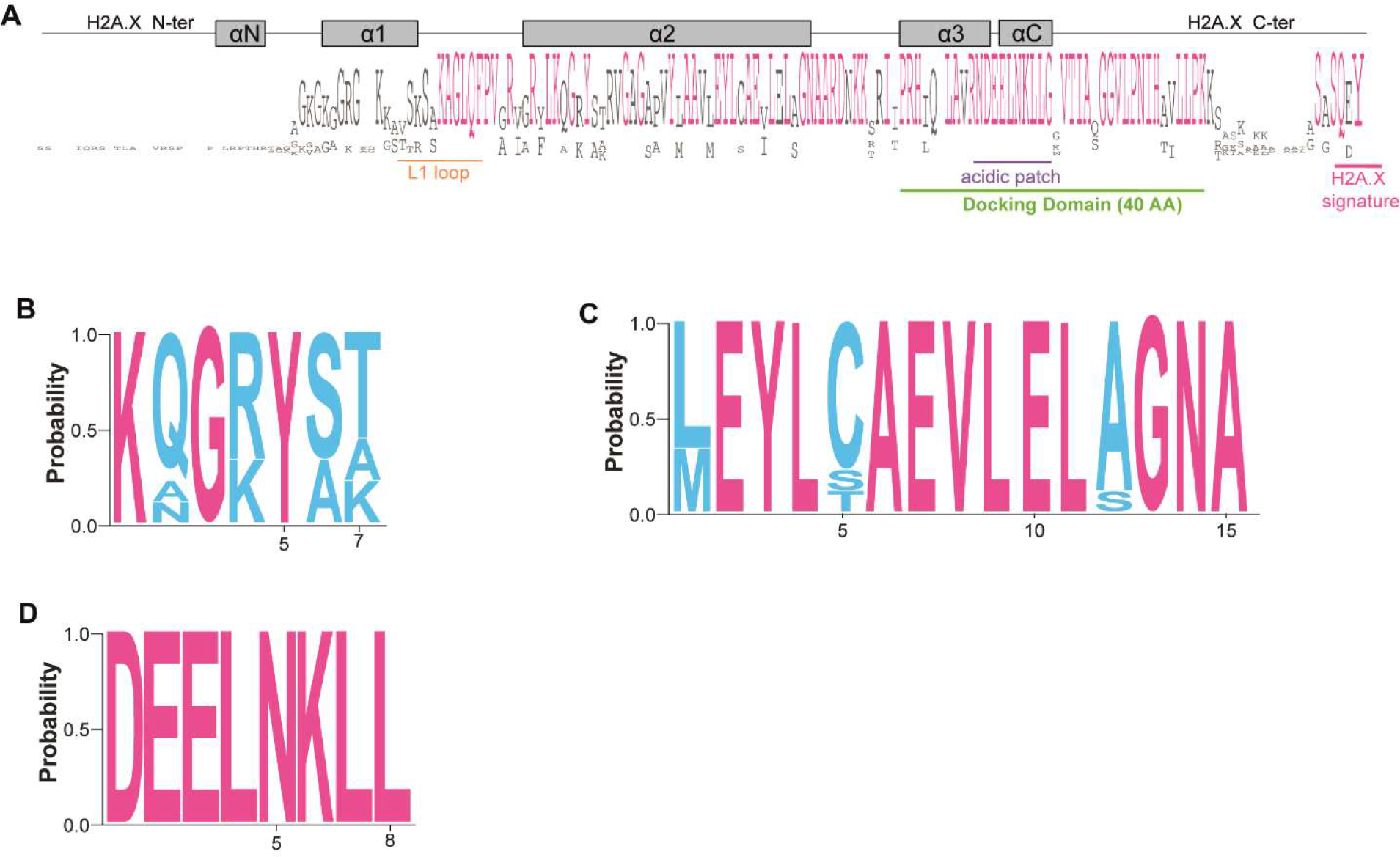
Analysis of the H2A.X variant in brown algae. *(A)* Consensus sequences for H2A.X proteins generated using Jalview (Waterhouse et al. 2009). The mature protein without the initial methionine is displayed. The L1 loop, acidic patch, docking domain and H2A.X signature are underlined in orange, purple, green and pink, respectively. The length of the docking domain is indicated (AA, amino acids). The helices are indicated by grey rectangles. *(B-D)* Logos of amino acid bias for the L1 loop, α2 helix and acidic patch.

### Identification of H2A variants with a H2A.Z signature in brown seaweeds

For each protein identified with a H2A.Z signature in the α2 helix (Jiang et al. 1998), we further analysed motifs present in the L1 loop, N- and C-terminal tails. This approach identified three types of proteins: (i) Group I with a long stretch of lysine and glycine in the N-terminal tail, a KSRVHSHQ motif in the L1 loop and a highly conserved TTKKRI motif in most C-terminal tails; (ii) Group II with a KYAT-related motif in the L1 loop motif; (iii) Group III with a long stretch of lysine and glycine in the N-terminal tail, a KYAT-related motif in the L1 loop motif and a highly conserved SQDY signature in the C-terminal tail (Fig. 3A). Based on the phylogenetic tree of H2A proteins (Fig. 3B), Group I proteins belonged to the monophyletic group of H2A.Z proteins. They also had the H2A.Z specific features: (i) a L1 loop with a single residue insertion (Fig. 4A-B); (ii) an acidic patch with an extra acidic residue (Fig. 4C) and (iii) a docking domain that was one amino acid shorter (Fig. 4A, Supplemental_Fig_S4). We thus named them H2A.Z and this variant was found as one single protein copy in all studied brown seaweed species.

**Figure 3:**
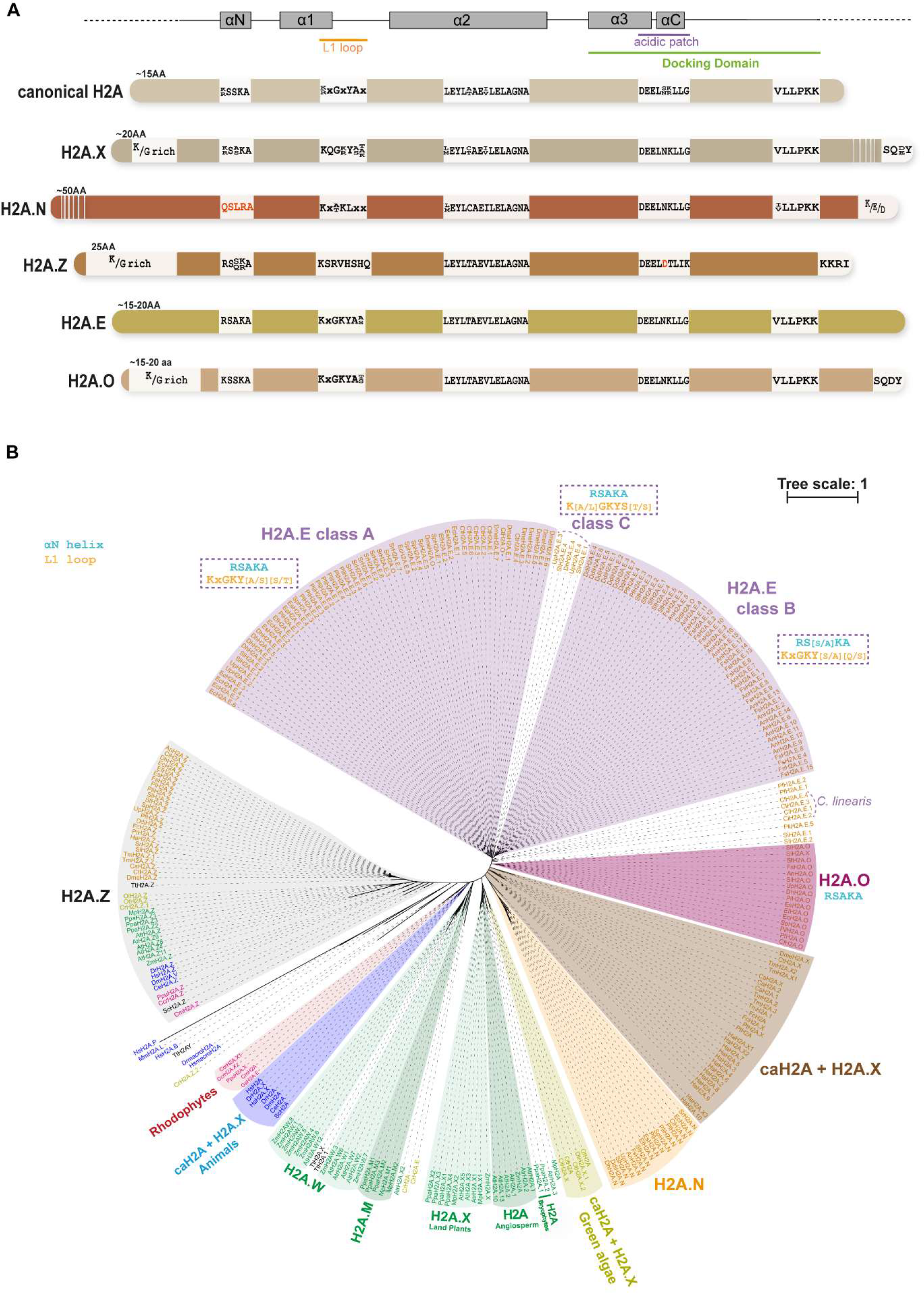
Identification of various H2A isoforms in brown seaweeds. *(A)* Schematic representation of canonical H2A and H2A variants identified in brown seaweeds. The consensus sequences for the αN helix, L1 loop, α2 helix and acidic patch are indicated. The motifs identified in the C-terminal tails of the H2A.X and H2A.Z and H2A.O variants are indicated. The presence of enriched amino acids (K/G rich) in N-terminal tails of H2A.X and H2A.Z and H2A.O variants is shown as well as in the C-terminal tail of the H2A.N variant (K/D/E). If present, the conserved sequence of the docking domain is indicated (mainly VLLPKK). The L1 loop, acidic patch and docking domain are underlined in orange, purple and green, respectively. The helices are indicated by grey rectangles. Black and red letters correspond to highly conserved or semi-conserved residues, respectively. The length of the N-terminal tail is indicated (AA, amino acids) for each H2A isoform as well as the length of the long C-terminal tail of the H2A.X variant. With the exception of the H2A.N N-terminal and H2A.X C-terminal tails, which are much longer than shown, boxes are proportional to the protein size. *(B)* Phylogenetic tree of H2A proteins. The H2A.E proteins group into two main classes, Class A and Class B, plus proteins from *C. linearis* that group together. To emphasize differences in proteins from various classes of variants, the consensus sequence is indicated in blue for the αN helix and in orange for the L1 loop. *Ascophyllum nodosum* (An), *Chordaria linearis* (Cl), *Choristocarpus tenellus* (Ct), *Chrysoparadoxa australica* (Ca), *Desmarestia herbacea* (Dh), *Dictyota dichotoma* (Ddi), *Discosporangium mesarthrocarpus* (Dme), *Ectocarpus crouaniorum* (Ec), *Ectocarpus fasciculatus* (Ef), *Ectocarpus siliculosus* (Es), *Fucus serratus* (Fse), *Heterosigma akashiwo* (Ha)*, Pleurocladia lacustris* (Pla), *Porterinema fluviatile* (Pf), *Pylaiella littoralis* (Pli), *Saccharina latissima* (Sl), *Sargassum fusiform* (Sf), *Schizocladia ischiensis* (Si), *Scytosiphon promiscuus* (Sp), *Sphacelaria rigidula* (Sri), *Tribonema minus* (Tm) and *Undaria pinnatifida* (Up) for brown algae. *Amborella trichopoda* (Atr)*, Arabidopsis thaliana* (At)*, Chlamydomonas reinhardtii* (Cr)*, Chondrus crispus* (Crr), *Cyanidioschyzon merolae* (Cm), *Danio rerio* (Dr)*, Drosophila melanogaster* (Dm)*, Fragilariopsis cylindrus (*Fc), *Galdieria sulphuraria* (Gs), *Homo sapiens* (Hs)*, Marchantia polymorpha* (Mp)*, Ostreococcus lucimarinus* (Ol), *Ostreococcus tauri* (Ot), *Phaeodactylum tricornutum* (Pt), *Physcomitrella patens* (Ppa)*, Porphyridium purpureum* (Ppu), *Saccharomyces cerevisiae* (Sc), *Tetrahymena thermophila* (Tt) and *Zea mays* (Zm).

**Figure 4:**
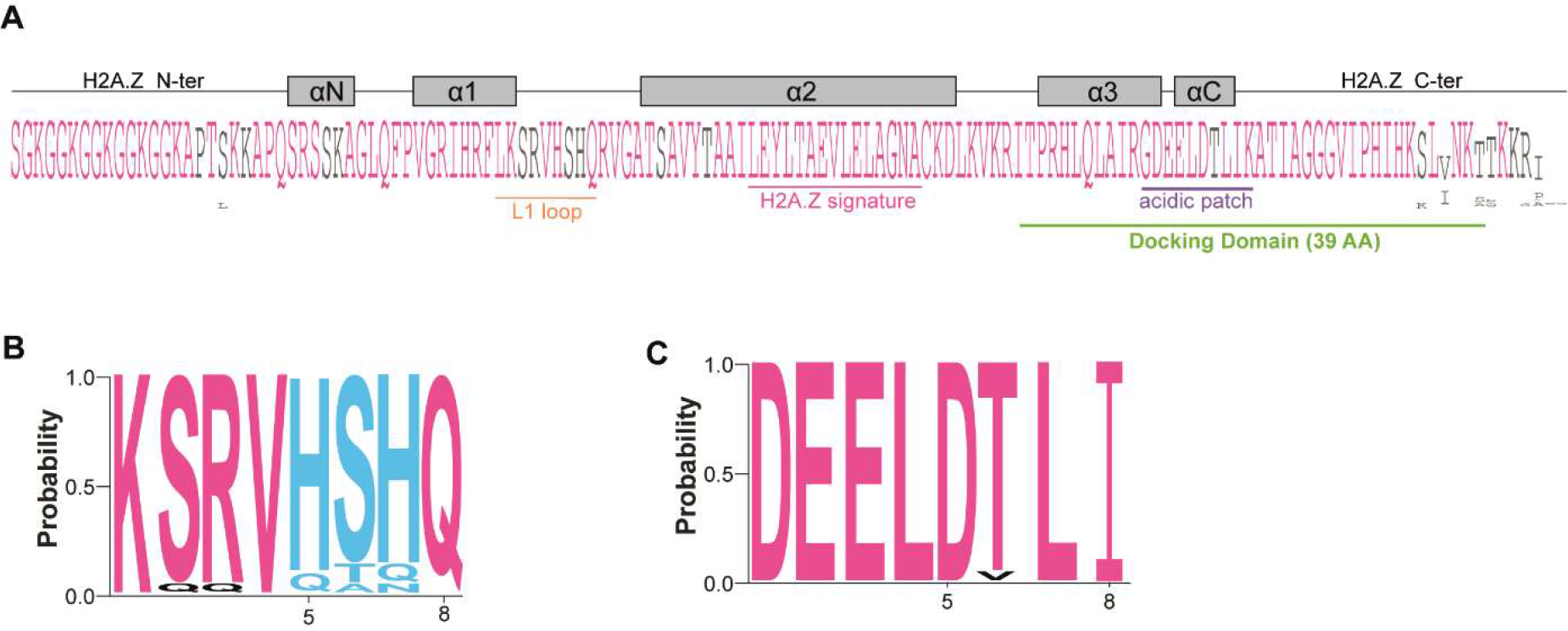
Analysis of the H2A.Z variant in brown algae. *(A)* Consensus sequences for H2A.Z proteins generated using Jalview (Waterhouse et al. 2009). The mature protein without the initial methionine is displayed. The L1 loop, acidic patch, docking domain and H2A.Z signature in the α2 helix are underlined in orange, purple, green and pink, respectively. The length of the docking domain is indicated (AA, amino acids). The helices are indicated by grey rectangles. *(B-D)* Logos of amino acid bias for the L1 loop and acidic patch.

Group II and III proteins are present in *S. ischiensis* and in the Phaeophyceae (Supplemental_Fig_S3). They have specific protein features and we named them H2A.E and H2A.O, following the histone nomenclature established by (Talbert et al. 2012). We defined motifs in their L1 loops as follows: KxGKY[A/S]x (Fig. 5B, H2A.E) and KxGKYA[T/S] (Fig. 6B, H2A.O). This motif appears to be a degenerated version of the brown algal H2A.X motif. Both H2A.E and H2A.O variants have a DEELNKLL motif in their acidic patch (Fig. 5C and 6C) and a docking domain of 40 amino acids ending with a VLLPKK sequence (Supplemental_Fig_S5A and Supplemental_Fig_S6A). Most H2A.O proteins clustered together phylogenetically, with a few exceptions that displayed a divergent αN helix sequence (Fig. 3B). The H2A.E proteins were organised into two main clades that differed by the αN helix and L1 loop consensus (Supplemental_Table_S2): (i) class A with RSAKA (Supplemental_Fig_S5B, left) and KxGKY[A/S] [S/T] (Supplemental_Fig_S5B, right) motifs, which contains half the H2A.E proteins; (ii) class B with RS[A/S]KA (Supplemental_Fig_S5C, left) and KxGKY[S/A][Q/S] (Supplemental_Fig_S5C, right) motifs. We also defined a class C, which constituted a third independent clade highly divergent in the L1 loop sequence (Supplemental_Fig_S5D, left). Additionally, H2A.E isoforms of *C. linearis* constituted an independent clade because of divergent αN helix and L1 loop (Supplemental_Fig_S5D) sequences. Therefore, the H2A.E and H2A.O variants constituted phylogenetically independent clades differing by the H2A.X C-terminal motif - harboured by H2A.O proteins - and by the number of isoforms, several for H2A.E and only one for H2A.O.

**Figure 5:**
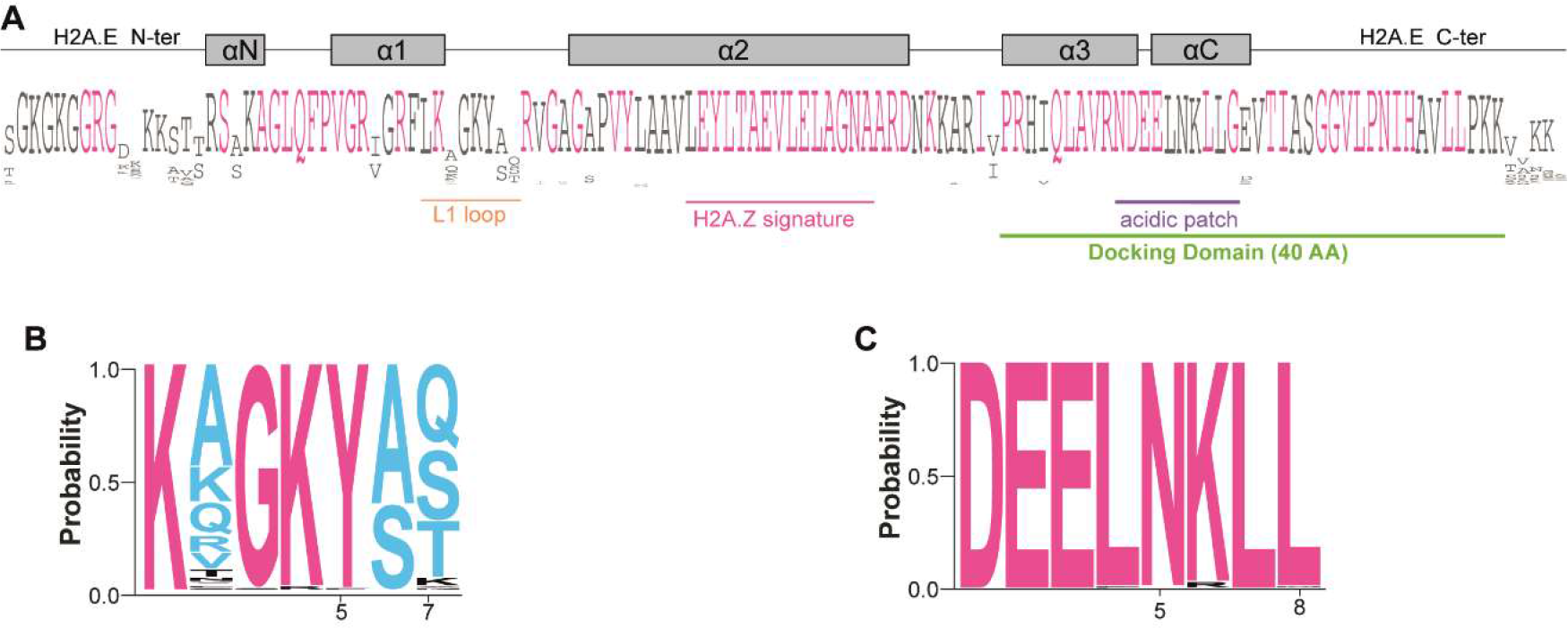
Analysis of the H2A.E variant in brown algae. *(A)* Consensus sequences for H2A.E proteins generated using Jalview (Waterhouse et al. 2009). The mature protein without the initial methionine is displayed. The L1 loop, acidic patch, docking domain and H2A.Z signature in the α2 helix are underlined in orange, purple, green and pink, respectively. The length of the docking domain is indicated (AA, amino acids). The helices are indicated by grey rectangles. *(B-D)* Logos of amino acid bias for the L1 loop and acidic patch.

**Figure 6:**
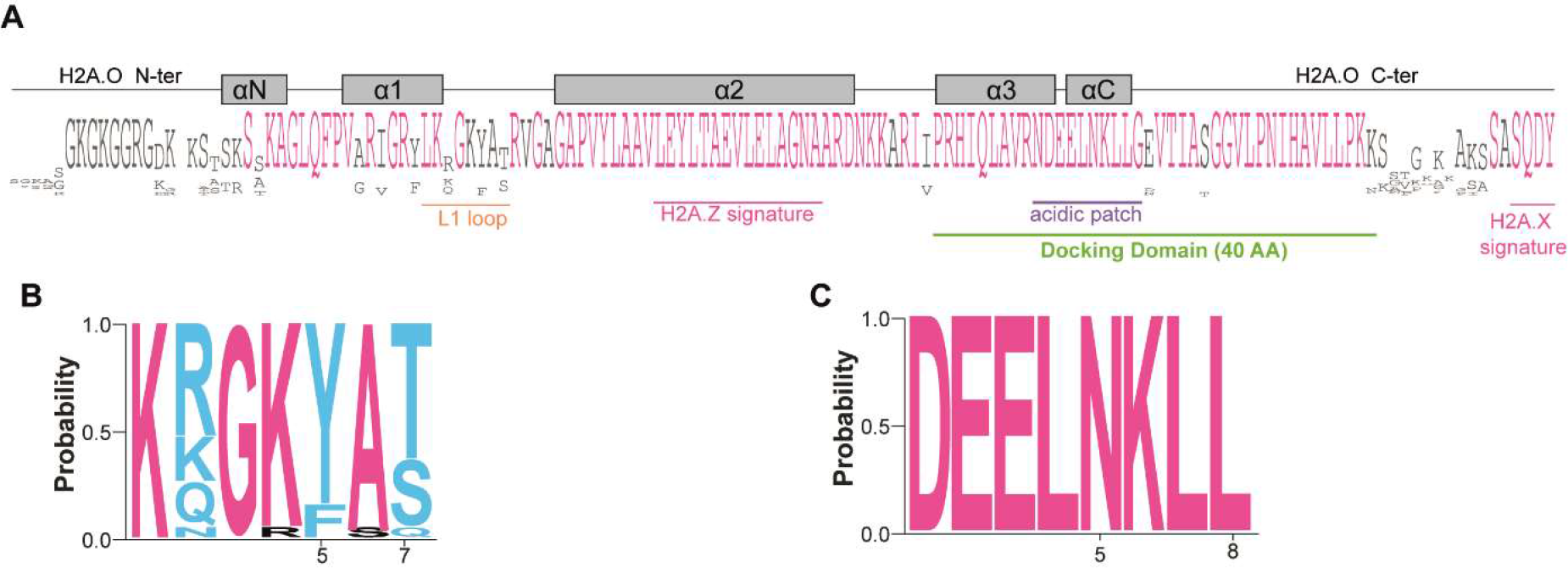
Analysis of the H2A.O variant in brown algae. *(A)* Consensus sequences for H2A.O proteins generated using Jalview (Waterhouse et al. 2009). The mature protein without the initial methionine is displayed. The L1 loop, acidic patch, docking domain and H2A.Z signature in the α2 helix as well as the H2A.X signature are underlined in orange, purple, green and pink, respectively. The length of the docking domain is indicated (AA, amino acids). The helices are indicated by grey rectangles. *(B-D)* Logos of amino acid bias for the L1 loop and acidic patch.

### The H2A.E variant was found in various algae species

In the Phaeoexplorer dataset, H2A.N, H2A.E and H2A.O variants were identified mainly in Phaeophycea (Supplemental_Fig_S3). This prompted us to explore whether such variants existed in other algae. Therefore, we searched for them in pennate diatoms (*Phaeodactylum tricornutum, Fragilariopsis cylindrus)*, unicellular green algae (*Ostreococcus tauri, Ostreococcus lucimarinus, Chlamydomonas reinhardtii*) and red algae (*Chondrus crispus, Galdieria sulphuraria, Porphyridium purpureum, Cyanidioschyzo merolae*) and in a recently released dataset of macroalgal deep genomics (Nelson et al. 2024). Both chosen diatoms possess canonical H2A, H2A.Z and H2A.X but lack H2A.N, H2A.E and H2A.O variants. Features of canonical H2A and H2A.X were similar between diatoms and brown seaweeds (Supplemental_Fig_S7A). Brown seaweed and diatom H2A.Z proteins had conserved features but diatom proteins lacked the TTKKRI C-terminal motif (Supplemental_Fig_S7B). Regarding green algae, both *Ostreococcus* species possess canonical H2A (Fig. 7A), H2A.X (Fig. 7B) and H2A.Z (Fig. 7C) but *C. reinhardtii* lacked a H2A.X variant (Rommelfanger et al. 2021) (Supplemental_Table_S1). The XP_001691545.1 protein from *C. reinhardtii*. has characteristic features of the H2A.E variant (*e.g.* KKGKYAE motif in its L1 loop and a H2A.Z signature in its α2 helix; Fig. 7D). Therefore, we named it CrH2A.E. This protein constituted a clade that was distinct from other H2A.E proteins and green algal H2A isoforms (Fig. 3B). We also identified an H2A.E in another class of Chlorophyta, in the macroalga *Avrainvillea amadelpha* (Fig. 7D). Regarding Rhodophyta, both *G. sulphuraria* and *C. merolae* lack an H2A.X variant. Based on the available data, the unicellular *G. sulphuraria* has only one H2A protein that has H2A.E-like signatures (*e.g.* a KNGNYAE motif in its L1 loop and a H2A.Z signature in its α2 helix; Fig. 7D). Hence, we named it GsH2A.E and it belongs to the Rhodophyta H2A clade (Fig. 3B). We also identified H2A.E proteins in another class of Rhodophyta, in the seaweed *Catenella fusiformis* (Fig. 7D), and a H2A.O variant in the Rhodophyta *Chroothece richterianum* (Fig. 7E). Therefore, we did not detect H2A.N variants in the green and red algal lineages but we did identify H2A.E and H2A.O variants in some green and red algae.

**Figure 7:**
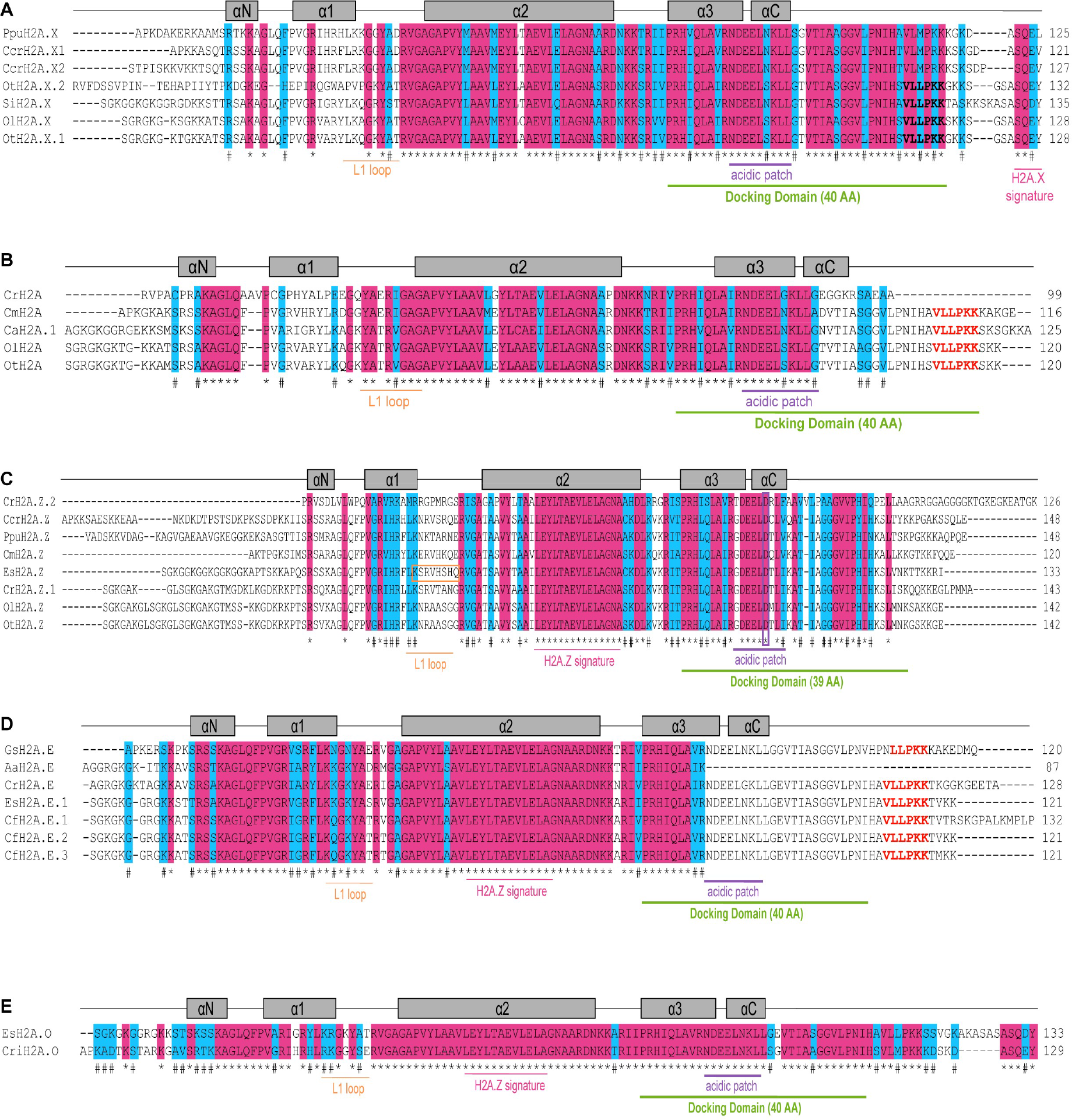
Analysis of the H2A isoforms in green and red algae. *(A-E)* Protein sequence alignment of the H2A.X variant (A), canonical H2A (B), H2A.Z (C), H2A.E (D) and H2A.O (E) variants. The L1 loop, acidic patch, docking domain and the H2A.X signature and the H2A.Z signature in the α2 helix are underlined in orange, purple, green and pink, respectively. The length of the docking domain is indicated (AA, amino acids). The helices are indicated by grey rectangles. Asterisks indicate a fully conserved residue and the hash conservation between residues with either strong or weak similar properties. *Chrysoparadoxa australica* (Ca), *Chlamydomonas reinhardtii* (Cr)*, Chondrus crispus* (Crr), *Cyanidioschyzon merolae* (Cm), *Ectocarpus siliculosus* (Es), *Galdieria sulphuraria* (Gs), *Ostreococcus lucimarinus* (Ol), *Ostreococcus tauri* (Ot), *Porphyridium purpureum* (Ppu), *Schizocladia ischiensis* (Si), *Catenella fusiformis* (Cf), *Avrainvillea amadelpha* (Aa), *Chroothece richterianum* (Cri).

### Identification of histone genes in the Phaeoexplorer genomes

Genes encoding canonical histones tend to be clustered while those coding for variants are more dispersed within animal genomes (Talbert and Henikoff 2010). Similarly, in *Ectocarpus* species 7 strain Ec32, canonical histones are organized into gene clusters (Bourdareau et al. 2021). In the present study, we identified the genes (Supplemental_Table_S3) encoding each H2A isoform and then analysed their genomic locations (Supplemental_Table_S4). We excluded from the analysis *Sargassum fusiform*, *Tribonema minus* and *Undaria pinnatifida* since their genome assemblies were of lower quality. The H2A.E variant has several isoforms (Supplemental_Table_S5) and each isoform was usually encoded by several genes (Supplemental_Table_S6) that were either dispersed or localised in tandem arrays or in close vicinity. Moreover, *H2A.E* genes co-localised with genes encoding H1, H2B, H3.1 and H4 (Supplemental_Table_S4), confirming the presence of histone gene clusters in brown seaweeds. Canonical H2A and the other H2A variants were encoded by single genes (Supplemental_Table_S3). Besides, canonical H2A genes were located on contigs that contain other histone-coding genes while *H2A.X*- and *H2A.O* genes were localised on contigs without such genes (Supplemental_Table_S4). Finally, we noticed that the *Ectocarpus siliculosus* male genome contained two tandem H2A.O-coding genes (contig73.15493.1 and contig73.15494.1, Supplemental_Table_S3). These findings indicate that brown seaweed genomes also have histone gene clusters.

### Selective deposition of PTMs on H2A variants

Histones are marked by a great variety of post-translational modifications. A mass spectrometry approach has been applied to *Ectocarpus* species 7 strain Ec32 to characterise histone PTMs (Bourdareau et al. 2021). We inspected the predicted peptides to assign these modifications to the corresponding H2A isoform(s) identified in the present study. The N-terminal tail of *Ectocarpus* species 7 H2A.Z is enriched in acetylated lysine residues (K3, K6, K9, K12, K15, K21) and has an acetylated S1 and a methylated R38 (Fig. 8A) (Bourdareau et al. 2021). The tail of diatom H2A.Z is also enriched in lysine residues and has been reported to be acetylated at S1/K3/K6/K9/K12/K15 but neither K21 acetylation nor R38 methylation was detected (Veluchamy et al. 2015). Both serine and lysine acetylation (S1, K3, K5) have been reported for another peptide (Bourdareau et al. 2021) that occurs in all the H2A.E isoforms and the H2A.O variant from *Ectocarpus* species 7 (Fig. 8B). We also investigated putative phosphorylation sites in the various H2A isoforms from *Ectocarpus* species 7, an aspect that was not investigated in the previous study (Bourdareau et al. 2021). The different *E. siliculosus* H2As variants have residues at several positions along the protein that could potentially undergo phosphorylation, with differences between variants (Supplemental_Fig_S8). Therefore, amino acid sequence differences between H2A variants potentially impact PTM deposition.

**Figure 8:**
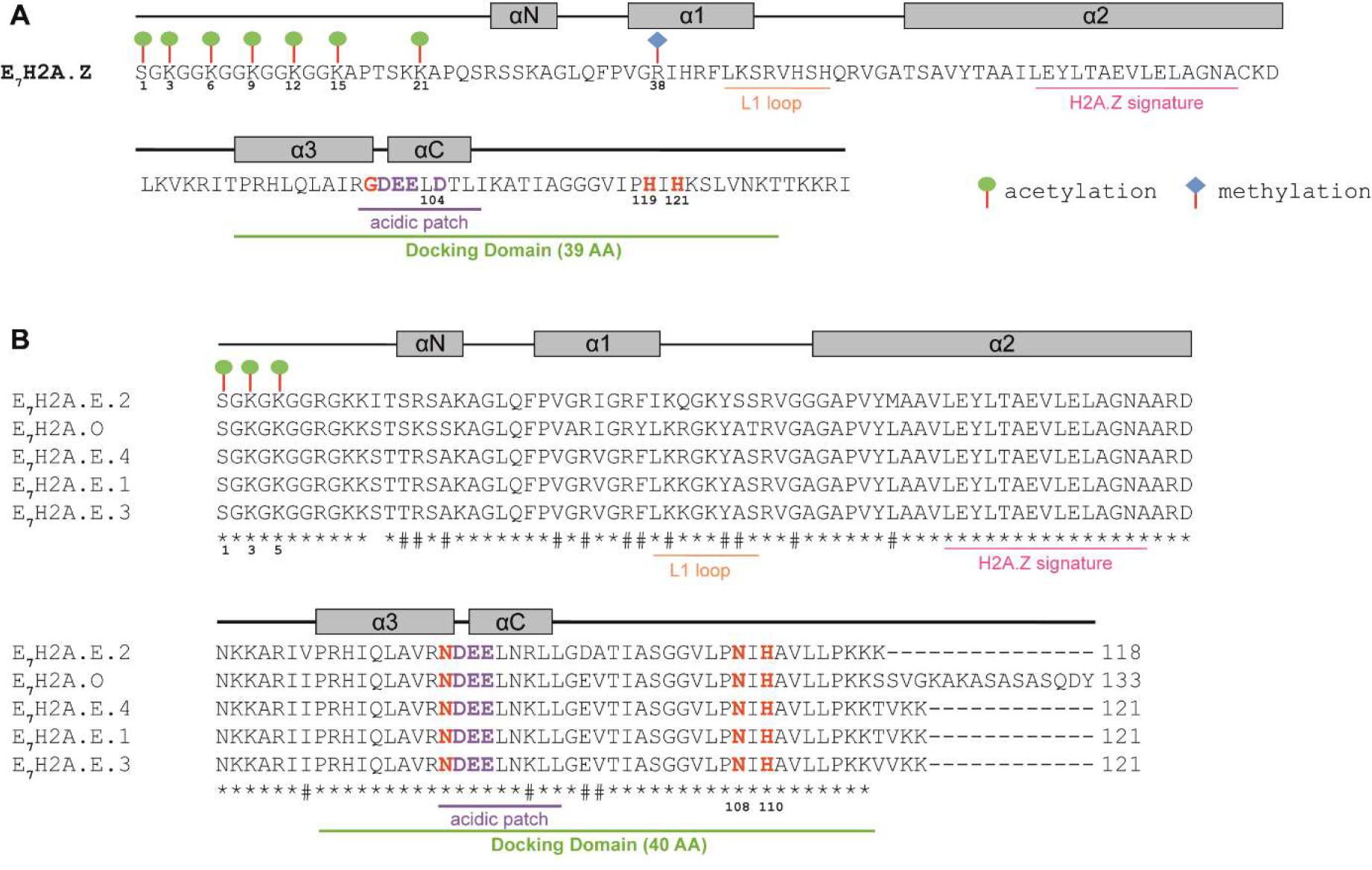
Predicted presence of PTMs on H2A isoforms from *Ectocarpus* species 7. *(A)* Protein sequence of the H2A.Z variant from *Ectocarpus* species 7. The two histidine residues present only in docking domains of H2A.Z proteins are displayed in red. Acidic patches start with a glycine in H2A.Z (displayed in red). The L1 loop, acidic patch, docking domain and the H2A.X signature and the H2A.Z signature in the α2 helix are underlined in orange, purple, green and pink, respectively. The length of the docking domain is indicated (AA, amino acids). The helices are indicated by grey rectangles. PTMs are shown based on data from (Bourdareau et al. 2021). *(B)* Alignment of *Ectocarpus* species 7H2A isoforms. In contrast to H2A.Z, the H2A.E and H2A.O variants harbor only one histidine residue (position 110, in red) in their docking domains, the first one is replaced by an asparagine (position 108, in red). Acidic patches start with an asparagine (displayed in red). Asterisks indicate a fully conserved residue and the hash conservation between residues with either strong or weak similar properties. PTMs are shown based on data from (Bourdareau et al. 2021).

### H2A variants were differentially expressed in reproductive tissues and in response to environmental conditions

Some histone variants have been reported to have sperm-specific expression patterns in *Arabidopsis* (Okada et al. 2005) or testis-specific expression patterns in mammals (Molaro et al. 2018). These variants can also be involved in development or stress responses (reviewed in Foroozani et al. 2022). We therefore evaluated expression of histone genes in various vegetative and reproductive tissues and under different growth conditions. Only *H2A*.Z and *H2A.N* genes were expressed in *Desmarestia herbacea* female and male gametophytes, with a greater transcript abundance in male compared to female (Supplemental_Fig_S9A). In *Dictyota dichotoma* (Supplemental_Fig_S9B), the *H2A.Z* gene was expressed with a similar pattern during the different life cycle stages (Supplemental_Fig_S9C). In this species, the *H2A.O* transcript abundance was high in female eggs, zygotes and embryos while those of *H2A.N* and *H2A.E* were differentially expressed throughout the life cycle with most genes being highly expressed in male sperm (Supplemental_Fig_S9C-D). In *Fucus serratus*, the *H2A.Z, H2A.N, H2A.O, H2A.E.3* and *H2A.E.6b* genes were expressed similarly in female and male gametophytes and in their gametes *i.e.* female eggs and male sperm (Fig. 9A). However, the *H2A.E.1* and *H2A.E.6a* genes were expressed only in male sperm, suggesting that H2A.E.1 could be a sperm-specific isoform. Analysis of *Ectocarpus* species 7 gametophyte strains (Gueno et al. 2022) showed that the *H2A.Z* gene was only expressed in females in contrast to the other seaweeds analysed where expression was also detected in males. However, the *H2A.N* and *H2A.O* transcript abundance was similar in male and female gametophytes (Supplemental_Fig_S9E).

**Figure 9:**
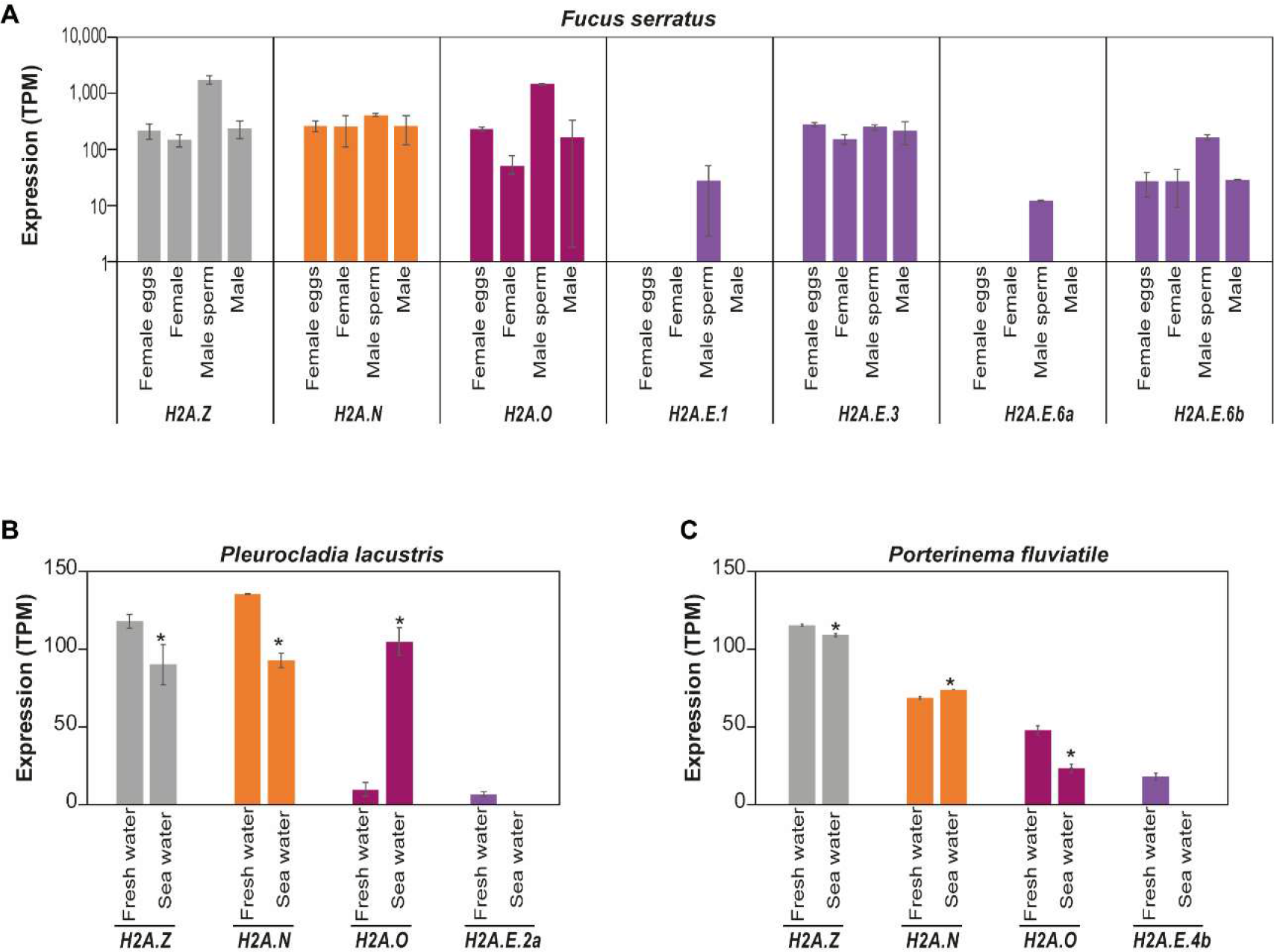
Analysis of the gene expression for the different H2A variants. *(A)* Expression of genes encoding *F. serratus* H2A variants. The histogram represents the mean transcript abundance in TPM (Transcripts Per Kilobase Million) displayed with a logarithmic scale. It corresponds to RNA-Seq data obtained from three biological replicates consisting of independent cultures of female and male gametophyte thalli (referred to as female and male, respectively) as well as from released female eggs and male sperm. Only genes with a TPM value above 2 are displayed. *(B-C)* Expression of the genes coding the *P. lacustris* (b) and *P. fluviatile* (c) H2A variants. The histogram represents the mean transcript abundance in TPM (Transcripts Per Kilobase Million). It corresponds to RNA-Seq data obtained from three biological replicates consisting of independent cultures grown either in fresh or sea-water. Only genes with a TPM value above 2 for at least one condition are displayed. Student’s t test; * P < 0.05.

To investigate if the diverse H2A variants could be involved in environmental responses, we compared gene expression in *Pleurocladia lacustris* and *Porterinema fluviatile* under two different experimental conditions: freshwater and seawater. In both species, the *H2A*.Z transcripts were more abundant in freshwater than in seawater and all the *H2A.E* genes were silenced under seawater conditions (Fig. 9B-C). However, the expression patterns of *H2A.N* and *H2A.O* differed in the two species under the freshwater and seawater conditions, suggesting that there are also species-specific differences (Fig. 9B-C). Interestingly, *P. lacustris H2A.E.2a* and *P. fluviatile H2A.E.4b* were the only *H2A.E* genes expressed in freshwater (Fig. 9B-C). Pairwise comparison of protein sequences revealed that encoded proteins were the most similar proteins (95.45% identity over 89% of the protein length) between the two species. Therefore, H2A variant genes were found to be differentially expressed depending on species, vegetative and reproductive tissues or environmental conditions, with the genes coding for H2A.E variants presenting the most variable expression patterns.

## Discussion

In this study, we carried out a comprehensive analysis to identify H2A isoforms in brown seaweed and sister taxa species, together with an analysis of available green and red algal genomes. Our study relied on a combination of seven features that were based on motif analysis (αN helix, L1 loop, α2 helix, acidic patch and C-terminal ending) and length (L1 loop and docking domain). This approach enabled us to identify the different variants present in brown seaweeds and to define their signatures (Fig. 3A). In addition to H2A.X and H2A.Z, which are both widespread eukaryotic variants, we report two novel variants present in three algal clades (Rhodophyta, Chlorophyta and Chromalveolata): the H2A.O and H2A.E variants. We also identified an additional variant, H2A.N, in most Phaeophyceae orders.

The H2A.E variant was not only found in brown seaweeds but also in two other independent lineages, Chlorophyta and Rhodophyta. Consequently, we hypothesize that the H2A.E variant evolved several times independently, as it has been proposed for canonical H2A and the H2A.X variant (Talbert et al. 2012). In addition, the number of H2A.E variants has expanded dramatically in brown seaweeds and this expansion may be linked to multicellularity rather than genome size. Indeed, *C. reinhardtii*, which has only one H2A.E protein, has a genome of similar size to some brown seaweeds. Moreover, in the red lineage, only the macroscopic seaweed *C. fusiformis* has three H2A.E proteins. Regarding H2A.O, this variant was found not only in the Phaeophyceae and Schizocladiophyceae but also in the red seaweed *C. richterianum*. Therefore, either this variant evolved at least twice in algae or brown algae acquired the gene through ancestral endosymbiosis of a red alga that already possessed the H2A.O variant. Under the second hypothesis, several Ochrophyta lineages would have then lost this variant during their evolutionary history. Concomitant with the acquisition of H2A.O, Phaeophyceae (with the exception of Discosporangiales) lost the H2A.X variants. Given this loss and based on features of the Discosporangiales H2A.X (*i.e.* a N-terminal tail with a long stretch of lysine and glycine and a C-terminal tail ending with a SQDY motif), the functions of H2A.X might be fulfilled by the H2A.O variant in species that have lost H2A.X. In addition, we observed that species that lack canonical H2A possess the H2A.N variant whose coding gene is expressed in most tissues and under most conditions (Fig. 9 and Supplemental_Fig_S9). This suggests that H2A.N could perform some roles of the canonical H2A, whose functional specialization has yet to be determined. The αN helices of all H2A.N variants shared a strongly divergent QSLRA motif, suggesting that the emergence of this specific helix was likely to be the first event that led to the appearance of the H2A.N variant. A H2A protein (KAJ3710985.1) from the fungus *Lentinula raphanica* has the same αN motif but without a long N-terminal tail. Moreover, the N-terminal tail of H2A.N has a PLRP motif (Supplemental_Fig_S1D, blue square). This motif was reported in the human E3 protein ligase and is characterized by a proline stack and a loop conformation catalytically critical for the function of this enzyme (Cappadocia et al. 2015). Therefore, the roles of the PLRP and QSLRA αN motifs remain to be investigated.

In the nucleosome, the H2A C-terminal tail interacts with DNA and protrudes from the nucleosome and this might enable interactions with various proteins and therefore lead to H2A specialization. Hence, we speculate that the H2A.Z and H2A.N C-terminal motifs (KKRI and K/E/D, respectively) could be involved in protein binding, in order to control chromatin remodelling for example. Moreover, the H2A.Z acidic patch begins with a glycine (Fig. 4) whereas it begins with an asparagine in all the other variants. This sequence variation could destabilize the H2A.Z/H3 interaction while the additional acidic residue in the docking domain might stabilize H2A.Z binding to the H4 N-terminal tail (Suto et al. 2000). Therefore, the different features observed in H2A variant sequences are likely to impact nucleosome stability and chromatin structure. The canonical H2A and the H2A.X, H2A.O and H2A.E variants are expected to behave similarly in terms of nucleosome stability since they have a similar length and L1 loop.

Histones are decorated by various post-translational modifications that modulate genome expression. The H2A.Z protein has several lysines in its C-terminal tail that are targets for acetylation (Fig. 8A) (Bourdareau et al. 2021). This hyper-acetylation might destabilize the nucleosome (Thambirajah et al. 2006). Post-translational modifications can also be involved in signalling. Indeed, during double-strand DNA break repair, serine and tyrosine residues in the H2A.X SQ[D/E]Y motif are phosphorylated, as is the upstream threonine residue (Li et al. 2010). Likewise, in this study, we reported the presence of one or two phosphorylatable residues in H2A.X (Fig. 2A) and H2A.O, respectively (Fig. 6A and Supplemental_Fig_S8), and the N-terminal tail of H2A.N was enriched in phosphorylatable serines (Supplemental_Fig_S8).

We investigated the tissue-specific expression of the various H2A variants throughout the complex life cycle of brown seaweeds as well as their regulation in response to environmental cues. The *H2A.Z, H2A.N* and *H2A.O* genes appeared to be ubiquitously expressed in most tissues and conditions for the analysed species. In contrast, almost all *H2A.E* genes were highly expressed in sperm cells of *F. serratus* and *D. dichotoma* (Fig. 9A and Supplemental_Fig_S9D). Female gametes *i.e.* eggs also expressed *H2A.E* genes but at a lower level, and not all of the genes were expressed (Fig. 9A and Supplemental_Fig_S9D). To remodel chromatin during mammalian spermatogenesis, testis-specific histones are incorporated into the chromatin and become hyperacetylated before being replaced by protamines. They are then exchanged for oocyte-derived histones after fertilization (reviewed in McLay and Clarke 2003). In plants, male gametophyte development is associated with the incorporation of specific histone variants, such as the *Arabidopsis* sperm-specific variant H3.10 (Okada et al. 2005). Moreover, pollen has specific histone post-translational modifications (reviewed in Borg and Berger 2015). Furthermore, gametogenesis is associated with marked changes in genome expression and is therefore associated with chromatin modifications such as DNA methylation (Feng et al. 2010). DNA methylation was not detected in *Ectocarpus* (Cock et al. 2010) but does occur in the diatom *P. tricornutum* (Veluchamy et al. 2013) and in *Saccharina japonica* but at a low level (Fan et al. 2020). Diatoms and most brown seaweeds lack orthologues of MET1/DNMT1 (DNA METhyltransferase 1 / DNA (cytosine-5)-MethylTransferase 1) (Maumus et al. 2011; Hoguin et al. 2023), the DNA methyltransferase involved in maintenance of CG methylation in plant and animal species. However, DNMT1 genes have been identified in *D. mesarthrocarpum* and two sister taxa of the brown algae *S. ischiensis* and *C. australica* (Denoeud et al. 2024). Therefore, DNMT1 appears to have been lost after divergence of the Discosporangiales from other brown seaweeds and concomitantly with the appearance of the H2A.N variant (Fig. 10). Given the absence of both DNA methylation and the H3K27me3 epigenetic mark (Bourdareau et al. 2021) in brown seaweeds, as well as the PRC2 complex responsible for writing this mark (Fig. 10), the H2A.E variant might have acquired specific functions to enable epigenetic reprogramming during Phaeophyceae oogenesis and spermatogenesis. For example, this could occur through post-translational modifications since the H2A.E variant contains a phosphorylatable serine at position 102 that is absent from the H2A.Z and H2A.N variants (Supplemental_Fig_S8). Epigenetic reprogramming is expected to occur in brown algae, probably through histone replacement and deposition of post-translational modifications. This study highlights the significance of understanding the impact of PTMs not only on canonical histones but also on specific histone variants, which may fulfil unique functions.

**Figure 10:**
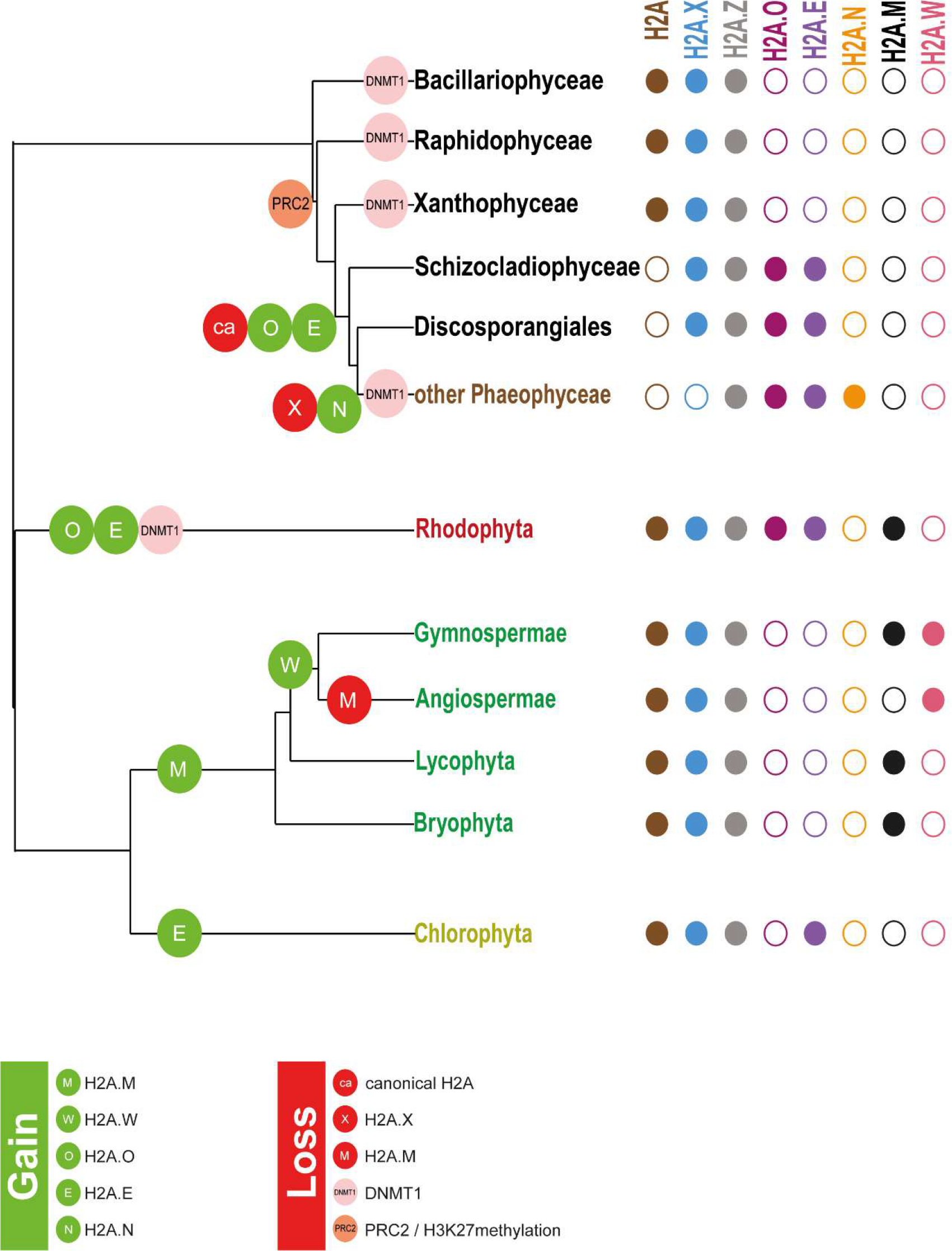
Evolution of some epigenetic features in photosynthetic organisms. The presented tree was constructed using TimeTree (Kumar et al. 2017). This figure displays the gain (green circles on tree branches) or loss (red, orange or pink circles on tree branches) of some epigenetic features (H2A variants, PRC2 complex/H3K27 methylation, DNMT1) in photosynthetic organisms (algae and land plants). Representative species were used for each clade: *Heterosigma akashiwo* (Raphidophyceae); *Tribonema minus* (Xantophyceae and Chrysoparadoxophyceae); *Schizocladia ischiensis* (Schizocladiophyceae); *Choristocarpus tenellus* (Discoporangiales); *Ectocarpus* (other Phaeophycea); *Phaeodactylum tricornutum* (Diatoms); *Cyanidioschyzon merolae*, *Chondrus crispus* and *Porphyridium purpureum* (Rhodophyta); *Chlamydomonas reinhardtii, Ostreococcus lucimarinus* and *Ostreococcus tauri* (Chlorophyta); *Marchantia polymorpha* (Bryophyta), *Selaginella moellendorffii* (Lycophyta); *Picea abies* (Gymnospermae); *Arabidopsis thaliana* (Angiospermae). DNA methylation was not detected in *Ectocarpus* (Cock et al. 2010). *Picea abies* has the DNMT1 homologues (Alakärppä et al. 2018), DNA methylation (Heer et al. 2018) and the epigenetic mark H3K27me3 (Nakamura et al. 2020). We performed a homology search for the catalytic subunit of the PRC2 complex that did not retrieved any homolog in sister clades of Phaeophyceae. Methylation of H3K27 was not detected in *Ectocarpus*, consistently with the absence of homologs for PRC2 core proteins (Bourdareau et al. 2021). In vascular plants, *Selaginella moellendorffii* has a PRC2 complex (Vijayanathan et al. 2022) and a DNMT1 homolog (Aliaga et al. 2019) and DNA methylation has been reported in *Ceratopteris richardii*, another vascular plant (McGrath and Pichersky 1997). For an easier display, we represented the gain of H2A.M based on the hypothesis that it evolved in land plants as H2A.W ancestor (Kawashima et al. 2015). On the right side of the tree, the presence or absence of an H2A isoform is depicted by a filled or unfilled circle, respectively. Each H2A isoform is depicted in a different colour with a code similar to that used in Supplemental_Fig_S3.

To conclude, brown seaweeds harbour a highly specific set of H2A variants, some of which are unique to this lineage and these might compensate for the loss of both DNA methylation and the PRC2 complex. The restricted distribution of these H2A variants among specific brown algal species suggests an adaptation to their various ecological niches and complex life cycles. In the future, it will be relevant to investigate the genomic localisation of the H2A variant proteins and, given the expanding applications of CRISPR-Cas9 mutagenesis (Badis et al. 2021), how loss of isoforms could impact chromatin structure and genome expression.

## METHODS

### Histone Identification in brown seaweed species using Phaeoexplorer genomic resources

Histone H2A protein sequence identification was performed by analysing: the 18 following brown seaweed species from the Phaeoexplorer genome resource (Denoeud et al. 2024) (https://phaeoexplorer.sb-roscoff.fr/home/): *Ascophyllum nodosum* (An), *Chordaria linearis* (Cl), *Choristocarpus tenellus* (Ct), *Desmarestia herbacea* (Dh), *Dictyota dichotoma* (Ddi), *Discosporangium mesarthrocarpus* (Dme), *Ectocarpus crouaniorum* (Ec), *Ectocarpus fasciculatus* (Ef), *Ectocarpus siliculosus* (Es), *Fucus serratus* (Fse), *Pleurocladia lacustris* (Pla), *Porterinema fluviatile* (Pf), *Pylaiella littoralis* (Pli), *Saccharina latissima* (Sl), *Sargassum fusiform* (Sf), *Scytosiphon promiscuus* (Sp), *Sphacelaria rigidula* (Sri) and *Undaria pinnatifida* (Up) and the four sister taxa species: *Chrysoparadoxa australica* (Ca), *Heterosigma akashiwo* (Ha)*, Schizocladia ischiensis* (Si) and *Tribonema minus* (Tm).

Histone H2A homologs were identified in these 22 species with the BLASTp tool run on the protein databases (https://blast.sb-roscoff.fr/phaeoexplorer/) using the canonical H2A protein sequence from the diatom *Phaeodactylum tricornutum*. To complete the analysis, genes and transcripts coding for the histones were retrieved from the genomes and predicted transcriptomes using BLAST (https://blast.sb-roscoff.fr/phaeoexplorer/). Proteins encoded by the identified genes and transcripts were predicted with the Expasy web tool (https://web.expasy.org/translate/) and validated with PANTHER (https://www.pantherdb.org/), Interproscan (Paysan-Lafosse et al. 2023) and Orthofinder (Emms and Kelly 2019). All the identified protein sequences and corresponding transcripts IDs are reported in Supplemental_Table_S3.

### Identification of histones in plants and animals

H2A protein sequences from *Homo sapiens* (Hs), *Mus musculus* (Mm), *Danio rerio* (Dr)*, Drosophila melanogaster* (Dm)*, Saccharomyces cerevisiae* (Sc), *Tetrahymena thermophila* (Tt) and *Zea mays* (Zm) correspond to protein sequences previously used in (Poulet et al. 2021). For *Chlamydomonas reinhardtii* (Cr)*, Chondrus crispus* (Crr), *Cyanidioschyzon merolae* (Cm), *Fragilariopsis cylindrus (*Fc), *Galdieria sulphuraria* (Gs), *Ostreococcus lucimarinus* (Ol), *Ostreococcus tauri* (Ot), *Phaeodactylum tricornutum* (Pt) and *Porphyridium purpureum* (Ppu), H2A protein sequences were retrieved from UniProt. For *Amborella trichopoda* (Atr)*, Arabidopsis thaliana* (At)*, Marchantia polymorpha* (Mp) and *Physcomitrella patens* (Ppa), the H2A protein sequences were described in (Kawashima et al. 2015).

### Identification of histones in algae from the green and red algal clades

*C. reinhardtii* predicted protein XP_001691545.1 is also referred as ch2a-IV / HTA2/HTA10 in (Khan et al. 2018) and H2A.0 in (Rommelfanger et al. 2021). Rommelfanger et al. (2021) predicted five proteins with minor sequence variations (H2A.0, H2A.1, H2A.2, H2A.3, H2A.4, Supplemental_Fig_S7C, divergent amino acids in red) and we thus speculated that they all correspond to XP_001691545.1. The A0A2K3DNT1 protein, referred in this study as CrH2A and as H2A.v2 in (Rommelfanger et al. 2021), seems to be incomplete based on its length (99 amino acids, Fig. 7B). The XP_001693700.1 and XP_001691141.1 proteins were referred as H2A.v and H2A.v3, respectively in (Rommelfanger et al. 2021). We used the name CrH2A to refer to A0A2K3DNT1 since it lacks any variant features (Fig. 7B). The XP_001693700.1 and XP_001691141.1 were named CrH2A.Z.1 and CrH2A.Z.2 since they have a L1 loop with an insertion and a H2A.Z signature in their α2 helix, as well as an acidic patch with one supplemental acidic residue and a docking domain of 39 amino acids (Fig. 7C). The XP_005537304.1 protein from the unicellular red alga *C. merolae,* lacks any variant signature and was named CmH2A. In this species, XP_005538662.1 has a H2A.Z signature in its α2 helix as well as one insertion in its L1 loop, an acidic patch with a supplemental acidic residue (DEEL**D**QLV) and a docking domain of 39 amino acids. XP_005538662.1was therefore named CmH2A.Z Supplemental_Table_S1). The red seaweed *C. crispus* and the unicellular red alga *P. purpureum* both possess H2A proteins with a SQ[E/D]Y-like motif without a H2A.Z signature in their α2 helix. These proteins (XP_005715904.1 and XP_005710906.1 and A0A5J4Z1G0) were named CrrH2A.X.1, CrrH2A.X.2 and PpuH2A.X, respectively (Supplemental_Table_S1). The two rhodophyta species *C. crispus* and *P. purpureum* also had H2A proteins with a H2A.Z signature in their α2 helix: XP_005715231.1 and A0A5J4YL73 that we named CrrH2A.Z and PpuH2A.Z. Both proteins have a L1 loop with an insertion typical of H2A.Z proteins, an acidic motif with a supplemental acidic residue and a docking domain of 39 residues (Fig. 7C). We identified H2A proteins with the BLAST tool run on the predicted protein database from (Nelson et al. 2024) (https://zenodo.org/records/7751045) using the H2A.Z protein sequence from *E. siliculosus*. We identified a H2A.E homologue in *A. amadelpha* with an incomplete sequence. However, genomic contigs for this species were not available at https://zenodo.org/records/7758534. We identified three H2A.E homologues in *C. fusiformis* and one H2A.O protein in *C. richterianum*. In this latter species, a tBLASTn search of genomic contigs did not identify any additional H2A proteins apart from the H2A.O variant.

### Phylogenetic analyses and protein sequence alignments

To generate the phylogenetic tree presented in Fig. 3B, selected sequences were first aligned in a multiple alignment with MUSCLE (Edgar 2004) using default parameters. Fast-Tree (v2.1.11) (Price et al 2021) was then applied with 1,000 bootstraps plus default parameters to generate the phylogenetic tree, which was drawn using the ITOL (Interactive Tree Of Life) tool (Letunic and Bork 2021). In the phylogenetic tree, plant, animal, green algal and red algal H2A proteins are displayed in dark green, blue, light green and pink. *P. tricornutum* proteins are depicted in red. For yeast and *T. thermophila*, they are displayed in black. The H2A proteins from brown seaweeds are displayed in brown. To compare the various H2A isoforms, multiple alignments were generated with Clustal Omega (Madeira et al. 2022).

### Analysis of consensus sequences

Consensus sequences were generated using Jalview Software (Waterhouse et al. 2009) and either global or local multiple alignments generated with Clustal Omega (Madeira et al. 2022). Logos were generated with the WebLogo web server (https://weblogo.threeplusone.com/) (Crooks et al 2004) using local multiple alignments. The amino acid height in the logo represents its relative frequency at its position.

### Identification of putative phosphorylation sites in H2A isoforms from *E. siliculosus*

Putative phosphorylation sites at serines, threonines and tyrosines residues of H2A isoforms (H2A.Z, H2A.N, H2A.O, H2A.E) from *E. siliculosus* were detected with NetPhos 3.1. Only scores higher than a specified threshold were retained as significant putative phosphorylation sites.

### Analysis of the expression for H2A variants

Differential expression data expressed in TPM (Transcripts Per Kilobase Million) were retrieved from RNA-Seq data generated for the following species: *F. serratus* (BioProject ID PRJNA731608)*, P. lacustris* (Denoeud et al. 2024)*, P. fluviatile* (Denoeud et al. 2024), *D. herbacea* (Denoeud et al. 2024), *D. dichotoma* (Bogaert et al. 2017). These data were obtained from the Phaeoexplorer project website (https://phaeoexplorer.sb-roscoff.fr/differential_expression/). For each species, all the RNA-seq data was mapped to the same reference genome and read counts generated using Salmon (version 1.3.0) (Patro et al. 2017). For *D. dichotoma,* samples were female and male gametophytes, eggs collected 15 min after release, sperm cells 1h after release, zygotes 1 h after release and embryos 8 h after fertilization (Bogaert et al. 2017) (Fig. Supplemental_Fig_S9B). For *F. serratus*, samples were female and male gametophyte thalli (referred to as female and male, respectively) as well as released female eggs and male sperm (Fig. 9A). For *Ectocarpus,* samples were collected from male (Ec457) and female (Ec460) *Ectocarpus* gametophyte lines and referred as female and male, respectively (Gueno et al. 2022). *D. herbacea, P. lacustris* and *P. fluviatile* samples were part of the project PRJEB72149. For *D. herbacea,* samples were female and male gametophytes grown in low light for one week before RNA extraction, cultures were transferred to high light to induce fertility (Denoeud et al. 2024). For the fresh water species *P. lacustris* strain SAG 25.93 and *P. fluviatile* strain SAG 2381, samples correspond to whole thalli grown in Petri dish cultures prepared with either diluted (5%) natural seawater or undiluted natural seawater (Denoeud et al. 2024).

## DATA ACCESS

Differential expression data were obtained from the Phaeoexplorer project website (https://phaeoexplorer.sb-roscoff.fr/differential_expression/). For *F.* serratus, samples correspond to the project PRJNA731608. *D. herbacea, P. lacustris* and *P. fluviatile* samples were part of the project PRJEB72149.

## ACKNOWLEDGMENTS

C.D. received a grant from the University of Nantes and the Région Pays de la Loire (M-EPICC, Etoiles Montantes). L.T. acknowledges support from the region of Pays de la Loire (ConnecTalent EPIALG project) and Epicycle ANR project (ANR-19-CE20-0028-02). This project was supported by the France Génomique National infrastructure project Phaeoexplorer (ANR-10-INBS-09).

## Author Contributions

C.D. and L.T. conceived and designed the study. J.M.A., J.M.C., F.D. and Z.N. provided genomic and transcriptomic resources. C.D. and E.R. analysed the data. C.D., J.M.C., E.R. and L.T. interpreted the data. C.D., J.M.C., E.R. and L.T. wrote the paper. J.M.A., F.D. and Z.N. contributed to the text editing.

